# From amazing work to I beg to differ - analysis of bioRxiv preprints that received one public comment till September 2019

**DOI:** 10.1101/2020.10.14.340083

**Authors:** Mario Malički, Joseph Costello, Juan Pablo Alperin, Lauren A. Maggio

**Affiliations:** Stanford University, Meta-Research Innovation Center at Stanford (METRICS); Uniformed Services University of the Health Sciences, Bethesda, Maryland, USA; Scholarly Communications Lab, Simon Fraser University, Vancouver, British Columbia, Canada; School of Publishing, Simon Fraser University, Vancouver, British Columbia, Canada

## Abstract

While early commenting on studies is seen as one of the advantages of preprints, the nature of such comments, and the people who post them, have not been systematically explored. We analysed comments posted between 21 May 2015 and 9 September 2019 for 1,983 bioRxiv preprints that received only one comment. Sixty-nine percent of comments were posted by non-authors (n=1,366), and 31% by preprint authors (n=617). Twelve percent of non-author comments (n=168) were full review reports traditionally found during journal review, while the rest most commonly contained praises (n=577, 42%), suggestions (n=399, 29%), or criticisms (n=226, 17%). Authors’ comments most commonly contained publication status updates (n=354, 57%), additional study information (n=158, 26%), or solicited feedback for the preprints (n=65, 11%). Our study points to the value of preprint commenting, but further studies are needed to determine the role that comments play in shaping preprint versions and eventual journal publications.

## Introduction

The practice of sharing preprint, authors’ versions of non-peer reviewed manuscripts, is on the rise. Once almost exclusively limited to the fields of high energy physics and economics on arXiv, RePec and SSRN preprint servers, preprints have gained much ground across a wide range of disciplines.^1–5^ Meta-research on preprints, however, remains scarce and mostly limited to the explorations of two servers: arXiv and bioRxiv. This limited research has shown that citation of preprints in scholarly literature had increased, and that articles first posted as preprints had higher citations rates and Altmetric scores than those not posted as preprints.^6–10^ Additionally, only minimal changes were found between preprints and the versions (of record) published in journals.^11–13^

In 2020, the COVID-19 pandemic led to a large increase in the posting of preprints, as well as scrutiny and the number of comments they receive on both social media platforms (e.g. Twitter) and comment sections of servers on which they are posted, with some comments prompting preprint retractions.^13, 14^ However, despite 70% of preprint servers allowing users to post comments on their platforms,^15^ and researchers perceiving the possibility of receiving comments as one of the advantages of preprints compared to traditional publishing,^16^ no research, to the best of our knowledge, has examined the nature of comments or actors involved in preprint commenting. In this study, which originated before the COVID-19 pandemic, we aimed to conduct an exploratory analysis of comments left on the bioRxiv servers. Furthermore, as at that time, the majority of preprints with comments, only a had single public comment, we decided to focus exclusively on such preprints.

## Methods

We conducted a cross-sectional study of bioRxiv preprints that received a single comment on the bioRxiv platform between 21 May 2015 (the earliest date available through the bioRxiv comment API) and September 9, 2019 (study data collection date).

### Data Collection

As part of our *Preprint Observatory* project,^17^ MM collected all available comments and related metadata using the bioRxiv comment API.^18^ Collected data included DOIs, links to the preprints, commenter username (i.e. proper name or created name), date and time the comment was posted, and the comment text. Data was stored and managed in Microsoft Excel. Initial extraction covered 6,454 comments posted to 3,265 unique preprints (which represents 6% of 56,427 preprints deposited on or before 9 September 2019; however, as the bioRxiv comment API did not provide access to comments posted before May 2015 this percentage does not represent the exact prevalence of commenting till September 2019). Of the 3,265 preprints with comments, 1,983 (60%) received only a single public comment, and we decided to focus on them in our first exploratory study. We enriched the data of those 1,983 comments by adding preprint authors, subject area classification, word count for comments, and published date of preprints as reported in Crossref and extracted during our *Preprints Uptake and Use Project*.^19^ Finally, we classified the commenters as authors or non-authors, and for authors we also captured their byline order (i.e. first, last or other – defined as neither first nor last).

### Data Analysis

Comments’ content classification was inductively derived using an iterative process of open-coding and constant comparison.^20^ Initial categories were devised by LAM based on a sample of 35 comments, and later expanded by MM using a sample of 200 comments. This initial categorization revealed distinct differences in content of comments left by authors and those left by non-authors.

### Identity of the commenter

MM and JC first checked whether each comment had been posted by an author of the preprint. This was done by comparing if the posted username matched any of the names of the preprint authors (and was helped by a simple full username search with any of the authors’ names - the simple search detected only 301 out of our later manually detected 617 cases as usernames often contained initials or symbols that were not an exact match with the names used in the preprint author byline). If the username was a pseudonym or a lab name, we classified the commenter as a non-author. During coding we amended our initial classification if the comments’ contents provided identification of the commenter.

### Content analysis

After grouping comments by the commenter type (author or non-author), MM, JC, and LM independently categorized all comments. Each comment could be classified to multiple categories. The only exception to this rule was if the comment was similar in structure and content to a full peer review report that is traditionally submitted as part of a journal peer review process. In those cases, we decided not to analyse the full contents of such review as they were often authored by multiple authors, contained multiple review reports, or included links to detailed reports posted on other websites. For all other comments, we classified the type of content they contained, but not the number of instances of each type they contained. For example, if the content type was a suggestion, we did not count the number of suggestions made in the comment, i.e. one suggestion for formatting a table, another for a figure, and additional suggestion for expanding the literature section. The three coders held weekly meetings online after coding batches of 200 to 300 comments. These meetings allowed for comparison of categorizations, resolving of differences, clarification of existing or introduction of new categories. Before each meeting, MM or JC would compare differences between the coders. If only one coder categorized a comment differently (e.g. did not mark a specific category) MM or JC re-read the comment, ruled on the found difference, and recorded the final categorization in the main database. When a single coder indicated a category the other two did not, or all coders disagreed on the categorization, the comment was marked and discussed at a weekly meeting until consensus was reached. We observed that our initial disagreement was most common for comments we categorised as suggestions or criticisms, and where tone, rather than content, dictated the categorisation (e.g. Comment 1: *“Great to see more well assembled lizard genomes, but it would have been nice to cite the more recent assemblies of…”;* Comment 2: *The authors state in the introduction that* [method] *has not been yet been reported". I beg to differ… following models have been generated and published*… [provides references to 3 studies]. We categorised comment 1 as suggestion, and comment 2 as criticism, based on their tone even though they both provided authors with additional references. As comments could have multiple categories, comment 1 was also classified as a praise).

While methods exist for calculating inter-rater reliability for data that could be classified as belonging to multiple categories,^21^ after each weekly meeting we only stored our agreed upon classification, so we cannot reconstruct the initial disagreements to produce such rating. It was also not our goal to study the difficulty of classifying comments, but rather, using a consensus approach, to explore the different types of comments posted on bioRxiv (before the pandemic). Our final classification tree and an example comment for each category are shown in Supplementary Table 1, and all comments and our assigned categories in our project’s database.^17^ Finally, to see if comments of preprints that received a single public comment, and that were the focus of our study, differed from first comments left for preprints which received more than one comment, we also randomly chose 200 of the latter preprints and analysed their first comments. This sub-analysis showed that all of these comments could be classified under our identified comment types.

### Statistical Analysis

We report absolute numbers and percentages for types of comments, and medians and interquartile ranges (IQR) for number of words per comment, number of comments per preprint and days from posting of the preprint to the comment. As number of words and days are integers, when medians or 25^th^ and 75^th^ percentiles had decimals, we rounded them to ease readability. Note on word count: As the texts of the comments were retrieved in HTML syntax, we replaced the hyperlink syntax (e.g. <a….a>) with the word LINK and counted it as only one word. When references were written out as *Author et al., year*, or *PMID: number*, those were counted by as many words as were written. Differences in number of words and time to publication between author and non-author comments were tested with Mann-Whitney test. We did not use time-to-event analysis as information for comments posted before May 2015 was not available through the API. Analysed comments came from all 27 bioRxiv subject area classifications (assigned by the authors during preprint upload, Supplementary Table 2), but despite there being slight differences in the number of comments per category; due to our sample size, focus on identifying comment types, and the fact that perceived preprint impact, as well as authors’ prestige, country and other factors might influence the posting of comments, we chose not to explore differences in commenting between subject areas. All analyses were conducted using JASP version 0.12.2.

## Results

Between 21 May 2016 and 9 September 2019, 1,983 bioRxiv preprints received a single public comment on the bioRxiv website. More than two thirds of those comments were posted by non-authors (n=1,366, 69%) while the remainder were posted by the preprint’s authors (n=617, 31%, Table 1). Overall, the non-author comments were longer than comments posted by the authors (median number of words was 38, IQR 17 to 83, for non-authors vs 18, IQR 11 to 32, for authors, Mann-Whitney test, P<0.001), and they were posted a median of 23 days (IQR 3 to 117) after the preprints. In comparison, authors’ comments were posted after a median of 91 days (IQR 3 to 23, Mann-Whitney test, P<0.001). Differences between types of comments with regards to number of words and days between preprint and comment publication are shown in Table 1.

**Table 1.** Type of comments, word count and days lapsed until comments were made on bioRxiv for preprints that received a single public comment between 21 May 2016 and 9 September 2019.

| Comment Type* | n (%)† | Median (IQR) no. of‡ |  |
| --- | --- | --- | --- |
|  |  | words | days<br>(from preprint to comment) |
| **Non-author's comment (n, total %)** | **1,366 (69)** | **38 (17 to 83)** | **23 (3 to 117)** |
| Praise | 577 (42) | 36 (17 to 70) | 13 (2 to 93) |
| Suggestion | 399 (29) | 46 (26 to 80) | 14 (2 to 90) |
| Criticism | 226 (17) | 78 (41 to 148) | 21 (3 to 104) |
| Asking for clarification | 213 (16) | 41 (22 to 73) | 31 (4 to 134) |
| Full peer review report | 168 (12) | 397 (78 to 785) | 46 (13 to 118) |
| Issue detected | 132 (10) | 22 (10 to 38) | 18 (2 to 80) |
| Asking for raw data or code | 41 (3) | 29 (13 to 52) | 21 (2 to 97) |
| Publication status update | 34 (2) | 12 (10 to 15) | 226 (96 to 343) |
| Other | 90 (7) | 20 (11 to 38) | 20 (3 to 128) |
| **Author's comment (n, total %)** | **617 (31)** | **18 (11 to 32)** | **91 (3 to 238)** |
| Publication status update | 354 (57) | 17 (10 to 20) | 178 (85 to 293) |
| Additional study information | 158 (26) | 21 (12 to 35) | 10 (2 to 121) |
| Soliciting feedback | 65 (11) | 21 (14 to 35) | 2 (1 to 8) |
| Study promotion | 44 (7) | 23 (14 to 63) | 1 (0 to 3) |
| Reply or thanks for received comments | 29 (5) | 45 (22 to 111) | 21 (5 to 98) |
| Other | 41 (7) | 28 (17 to 48) | 2 (1 to 17) |

\* Examples of comment types can be found in Supplementary Table 1.
† Percentages do not add up to a 100, as comment's content could contain more than one comment type.
‡ The median and IQRs for number of words in this table are calculated based on the total number of words of a comment, not its individual parts (i.e. if a comment contained both a praise and a suggestion, its total word count is added to both of those categories).

Twelve percent of non-author’s comments (n=168) were full review reports resembling those traditionally submitted during the journal peer review process. They were authored by either single individuals (n=87, 52%) or group of authors (n=81, 48%, Supplementary Table 3). The latter most commonly published their review following a journal club discussion (n=41, 51%). Comments not resembling full peer review reports most commonly praised the preprint (n=577, 42%), made suggestions on how to improve it (n=399, 29%), or criticized some aspect of the preprint (n=226, 17%, Table 1). Praise was most commonly found alongside suggestions (n=201; 50%) or comments asking for clarifications (n=101; 47%), and least commonly alongside comments that criticised the preprint (n=70, 31%), reported issues (n=38, 29%) or that inquired of the preprints publications status (n=9, 26%, Supplementary Table 4). Praise words alone (e.g. “*Amazing work!”*) constituted 86 (6%) comments. Comments containing suggestions (n=399, 29%) often included suggestions of literature (co-)authored by the commenter (n=143, 36%) or suggestions of other literature (n=118, 9%, Supplementary Table 3).

Lastly, we present some examples of the comments we classified as belonging to the "other” category (a full list of those comments available on our project website). There were three comments that raised research integrity issues (a possible figure duplication, an undeclared conflict of interest, and use of bots to inflate paper download numbers). There were also comments that raised personal issues. In one comment a parent requested more information on a rare disease (covered by the preprint) that was affecting their children, and in another case an individual inquired about possible PhD mentors for a topic related to the preprint. There were also comments that touched upon the culture of preprinting, with one comment asking authors to include brief summaries of what had changed between preprint versions, another expressing a view that preprints make traditional publishing redundant, and one praising authors for replying to questions they asked through email. Similarly, one comment we classified as full peer review report, also included a statement of hope “*to get more comments on bioRxiv*…*prior to submission to a peer reviewed-journal”* as they would “*rather have a revised pre-print than a correction / retraction” in a journal*.

Authors’ comments most commonly contained updates about the preprint’s publication status (n=354, 57%), additional information on the study (n=158, 26%), or solicited feedback for the preprint (n=65, 11%, Table 1). Of all authors’ comments, 321 (52%) were posted by the first author of the preprint, 209 (34%) by the last, and 65 (11%) by other authors (we could not identify the byline order for four percent of comments, n=22, as the registered username was either a pseudonym, e.g. *W1ndy*, or a lab name, e.g. *Lewy Body Lab*). A small percentage (n=29, 5%) of author comments were replies to feedback authors received elsewhere, e.g. during peer review or through personal emails (Supplementary Table 4). Lastly, as above, we present few examples of authors’ comments classified as belonging to the “other” category (with full list of those available on our project website). In five comments authors requested suggestions on where to publish their preprint, and in one comment authors mentioned that an editor saw their preprint and invited them to submit it to their journal. In one comment, an author alerted the readers of an error in a figure and also playfully chided (using a smiley emoticon) the co-author for hastily uploading the files before checking them. In another co-authors alerted readers that the preprint had been posted without the approval of the co-authors and urging the scientific community to ignore this version (to date the preprint in question has not been retracted).^22^ Finally, in one example (of a comment classified as a publication status update), the author said they did not plan to submit the preprint to a journal, as publishing on bioRxiv makes it freely available to everyone.

## Discussion

Our study of preprints posted on the bioRxiv server that received a single public comment found that more than two thirds of those comments were left by non-authors and most commonly praised, offered suggestions, or criticised the preprints. Additionally, almost a sixth of these non-author comments contained detailed peer review reports akin to those traditionally submitted during the journal peer review process. These findings support previous studies that showed the opportunity to receive feedback was perceived as one the benefits of preprints compared to traditional publishing.^23^ However, we also found that less than ten percent of all bioRxiv preprints received public comments before the COVID-19 pandemic. This low prevalence of scholarly public commenting has been previously observed for post-publication commenting of biomedical articles, and was the reason for discontinuing PubMed Commons, the National Library of Medicine’s commenting resource.^24^ Similar low prevalence of post-publication commenting has also been found across disciplines on PubPeer.^25^ Nevertheless, as has been stated for those services,^24, 25^ some of the comments have been crucial for scholarly debates and even led to retractions of papers, a practice also observed for bioRxiv preprints.^26^ In our study, we observed that eleven percent of authors’ comments were actively inviting others to comment on their preprint, with one comment explicitly stating that they would rather make changes to the preprint than to a version published in a journal.

The lack of traditional peer-review is often perceived as the biggest criticism of pre-printing, alongside cases of information misuse^27^ and posting of low-quality studies. Thus, bioRxiv (alongside arXiv and medRxiv) have displayed clear disclaimers for COVID-19 preprints that state preprints are “*preliminary reports that have not been peer-reviewed”* and they should not be “*reported in media as established information*”.^28^ Related to this criticism and the benefits of preprint commenting, there has also been a rise of specialised preprint review services (e.g. [PreReview](https://prereview.org/),^29^ [ReviewCommons](https://www.reviewcommons.org/),^30^ [PeerageofScience](https://www.peerageofscience.org/))^31^ or overlay journals (e.g. [RapidReviews](https://rapidreviewscovid19.mitpress.mit.edu/),^32^ [DiscreetAnalysis](https://discreteanalysisjournal.com/))^33^ aimed at providing expert reviews for preprints, or endorsement of preprints (e.g. [Plaudit](https://plaudit.pub/)).^34^ On a similar note to emphasize the possible role that commenting has in the scientific discourse, reference software Zotero can display references that have PubPeer comments,^35^ and a recently launched biomedical search engine [PubliBee](https://www.publibee.com/#/search),^36^ implemented (up)voting of comments.

Alongside posting of full peer review reports, our study also confirmed other known practices and potential benefits associated with the preprinting culture. For example, using preprints as final publication outputs, soliciting or being invited by editors to publish studies posted as preprints, calling out suspected research integrity issues, engaging in discussion or proposing collaborations, as well as publishing of peer review reports from those training on how to conduct peer review or from journal club discussions. These findings may provide authors encouragement to consider or continue depositing preprints.

Furthermore, we have shown that almost a third of the comments were left by the authors of the preprints, and their comments were mostly updates of preprints’ publication status or additional information about the studies. Authors’ comments were also in general left after a much longer period than those of the non-authors. This aligns with found median times of 166 to 182 days between posting a preprint on bioRxiv and publication of that study in a journal,^37,38^ which were similar to the median time of 172 days we found for comments on publication status updates.

### Limitations

We did not attempt to define if non-authors that posted comments were indeed peers, nor did we compare their expertise or publication records with those of the authors of the preprint on which they were commenting. We are also aware that some comments were left by patients, students and the individuals that stated a lack of expertise in the field. However, defining and soliciting feedback from a competent peer is known to be difficult,^39^ with previous studies demonstrating minimal agreements between peers assessing the same study.^40^ Furthermore, we did not attempt to define the quality of the comments, nor if the contents of comments (e.g. raised criticisms or suggestions) were indeed valid. We also did not check if comments led to changes or updates of the preprints or eventual published manuscripts, nor if the authors were even aware of them. Regarding the latter, as we analysed preprints that only had a single comment, none of the authors used the preprint platform to reply to them. We however did find that five percent of authors’ comments were replies to comments or peer reviews they received elsewhere, and we did encounter an example of a non-author comment that indicated they communicated with the authors by email. The purpose of our research was not to provide external validity of the claims stated in the comments, but rather showcase, for the first time, the most common types of comments left on the platform (before the COVID-19 pandemic). Our study is also limited in that we did not analyse discourse that might occur in preprints which received multiple comments. However, we did analyse the first comments of a random sample of 200 of such preprints to confirm that they do fall within the categories analysed here. Finally, we acknowledge that our backgrounds are not in biology, and that this may have affected our ability to make a clear distinction between some comment types, especially in distinguishing between suggestions and criticisms. We however feel that the observed differences in the number of words between our identified comment types, as well as prevalence or praise which is more common for comments that contained suggestions than criticisms, provides support for our categorization.

### Future Directions

In the last year, the landscape of scientific communication has been significantly altered by the COVID-19 pandemic. Looking ahead, in our future research, we will examine comments and exchanges that occur when multiple comments are posted for the same preprint, as well as the possible difference in types of comments during the COVID-19 pandemic. If warranted, additional research and discourse analysis should be conducted to better understand preprint commenting, as well as the factors associated with authors willingness to respond to comments or incorporate feedback into preprint updates or submitted manuscripts for publication. Nevertheless, as we have previously advocated describing changes between preprint versions, as well as between preprints and published versions of record,^41^ perhaps it is also time for commenting sections and social media platforms to implement categorization of comments and even quality evaluation or upvoting of comments.^42^

### Conclusions

Despite a small prevalence of commenting occurring on bioRxiv before the COVID-19 pandemic, the non-author comments contained many aspects of feedback seen in traditional peer review, while those by the authors most commonly addressed the publication status of the preprints or provided additional information on the study. Based on the types of comments we identified, bioRxiv commenting platform appears to have potential benefits for both the public and the scholarly community. Further research could measure the direct impact of these comments on later preprint versions or journal publications, as well as the feasibility and sustainability of maintaining and moderating commenting sections of bioRxiv or other preprint servers. Finally, we believe that user-friendly integration of comments from server platforms and those posted on social media (e.g. Twitter) and specialized review platforms would be beneficial for a wide variety of stakeholders, including the public authors, commenters, and researchers interested in analysis of comments.

## Acknowledgments

This project was conceived and inspired during the collaboration of [ASAPbio](https://asapbio.org/) and [ScholCommLab](https://www.scholcommlab.ca/) on the [PreprintsUptakeandUseProject](https://www.scholcommlab.ca/research/preprints/). Our preliminary analysis was presented at the [2ndInternationalConferenceonPeerReview](https://www.peere.org/conference/). We would like to thank, in alphabetical order, Cathrine Axfors, Lex M. Bouter, Noah Haber, Ana Jerončić, and Gerben ter Riet for their comments on the draft of our manuscript.

## Notes

**Funding:** Elsevier funding was awarded to Stanford University for a METRICS postdoc position that supported MM’s work on the project.

https://data.mendeley.com/datasets/zrtfry5fsd/4

### Competing Interest Statement

The authors have declared no competing interest.

## References

1. Cobb M. The prehistory of biology preprints: A forgotten experiment from the 1960s. PLOS Biology. 2017;15(11):e2003995.

2. Rittman M. Preprint Servers. 2018. Available from: http://researchpreprints.com/.

3. Tomaiuolo NG, Packer JG. Pre-print servers: pushing the envelope of electronic scholarly publishing. Searcher. 2000;8(9):53.

4. Lin J. Preprints growth rate ten times higher than journal articles: Crossref.org; 2018 updated September 11 2018. Available from: http://www.crossref.org/blog/preprints-growth-rate-ten-times-higher-than-journal-articles/.

5. Anaya J. Monthly Statistics 2018. Available from: http://www.prepubmed.org.

6. Serghiou S, Ioannidis JP. Altmetric Scores, Citations, and Publication of Studies Posted as Preprints. JAMA. 2018;319(4):402–4.

7. Li X, Thelwall M, Kousha K. The role of arXiv, RePEc, SSRN and PMC in formal scholarly communication. Aslib journal of information management. 2015;67(6):614–35.

8. Feldman S, Lo K, Ammar W. Citation Count Analysis for Papers with Preprints. arXiv preprint arXiv:180505238. 2018.

9. Larivière V, Sugimoto CR, Macaluso B, Milojević S, Cronin B, Thelwall M. arXiv E-prints and the journal of record: An analysis of roles and relationships. Journal of the Association for Information Science and Technology. 2014;65(6):1157–69.

10. Alvarez GR, Caregnato SE. Preprints na comunicação científica da Física de Altas Energias: análise das submissões no repositório arXiv (2010-2015). Perspectivas em Ciência da Informação. 2017;22:104–17.

11. Klein M, Broadwell P, Farb SE, Grappone T. Comparing published scientific journal articles to their pre-print versions. International Journal on Digital Libraries. 2019;20(4):335–50.

12. Carneiro CFD, Queiroz VGS, Moulin TC, Carvalho CAM, Haas CB, Rayêe D, et al. Comparing quality of reporting between preprints and peer-reviewed articles in the biomedical literature. bioRxiv. 2020:581892.

13. Fraser N, Brierley L, Dey G, Polka JK, Pálfy M, Coates JA. Preprinting a pandemic: the role of preprints in the COVID-19 pandemic. bioRxiv. 2020.

14. Fang Z, Costas R. Tracking the Twitter attention around the research efforts on the COVID-19 pandemic. arXiv preprint arXiv:200605783. 2020.

15. Malički M, Jeroncic A, ter Riet G, Bouter LM, Ioannidis JP, Goodman SN, et al. Preprint servers’ recommendations for transparency and research integrity: a cross-sectional study across scientific disciplines. 2020.

16. Chiarelli A, Johnson R, Pinfield S, Richens E. Preprints and Scholarly Communication: Adoption, Practices, Drivers and Barriers [version 1; peer review: 2 approved with reservations]. F1000Research. 2019;8(971).

17. Malički M. Preprint Observatory. 2020. http://data.mendeley.com/datasets/zrtfry5fsd/4.

18. bioRxiv. bioRxiv API (beta). Disqus Comments. API Summary. Available from: http://connect.biorxiv.org/api/disqus/help.

19. Malicki M, Alperin JP. Preprints Uptake and Use Project 2019. Available from: http://www.scholcommlab.ca/research/preprints/.

20. Straus A, Corbin J. Basics of qualitative research: Grounded theory procedures and techniques: Newbury Park, CA: Sage; 1990.

21. Janson H, Olsson U. A measure of agreement for interval or nominal multivariate observations. Educational and Psychological Measurement. 2001;61(2):277–89.

22. Narayanan V, Utomo WK, Bruno MJ, Peppelenbosch MP, Konstantinov SR. Bacterial invasion of the pancreas revealed after analyses of the pancreatic cyst fluids. bioRxiv. 2016:064550.

23. Chiarelli A, Johnson R, Richens E, Pinfield S. Accelerating scholarly communication: the transformative role of preprints: Zenodo; 2019.

24. NCBI Insights Internet2018. Available from: http://ncbiinsights.ncbi.nlm.nih.gov/2018/02/01/pubmed-commons-to-be-discontinued/.

25. McCook A. Retraction Watch Internet2018. Available from: http://retractionwatch.com/2018/02/02/pubmed-shuts-comments-feature-pubmed-commons/.

26. Oransky I, Marcus A. Quick retraction of a faulty coronavirus paper was a good moment for science. STAT. 2020.

27. Gallotti R, Valle F, Castaldo N, Sacco P, De Domenico M. Assessing the risks of“ infodemics” in response to COVID-19 epidemics. arXiv preprint arXiv:200403997. 2020.

28. COVID-19 SARS-CoV-2 preprints from medRxiv and bioRxiv 2020. Available from: http://connect.biorxiv.org/relate/content/181.

29. 2017. Available from: http://elifesciences.org/labs/57d6b284/prereview-a-new-resource-for-the-collaborative-review-of-preprints.

30. 2019. Available from: http://www.embo.org/news/press-releases/2019/review-commons-a-pre-journal-portable-review-platform.html.

31. Seppänen J-T. 2016. Available from: http://blogs.biomedcentral.com/bmcblog/2016/06/16/peerage-science-inspiration-aims-future-developments/.

32. Heidt A. 2020. Available from: http://www.the-scientist.com/news-opinion/new-journal-to-publish-reviews-of-covid-19-preprints-67675.

33. Lobo MP. A do-it-yourself overlay journal. 2019.

34. Pfeiffer N. 2019. Available from: http://www.cos.io/blog/now-you-can-endorse-papers-osf-preprints-plaudit#:~:text=To%20endorse%20a%20preprint%2C%20start,clear%2C%20or%203)%20exciting.

35. PubPeer Extensions 2020. Available from: http://pubpeer.com/static/extensions.

36. Publibee 2020. Available from: http://www.publibee.com/#/search.

37. Abdill RJ, Blekhman R. Meta-Research: Tracking the popularity and outcomes of all bioRxiv preprints. eLife. 2019;8:e45133.

38. Fu DY, Hughey JJ. Releasing a preprint is associated with more attention and citations. bioRxiv. 2019:699652.

39. Glonti K, Cauchi D, Cobo E, Boutron I, Moher D, Hren D. A scoping review on the roles and tasks of peer reviewers in the manuscript review process in biomedical journals. BMC medicine. 2019;17(1):118.

40. Bornmann L, Mutz R, Daniel H-D. A reliability-generalization study of journal peer reviews: A multilevel meta-analysis of inter-rater reliability and its determinants. PloS one. 2010;5(12):e14331.

41. Malički M, Alperin JP. 2020. Available from: http://www.scholcommlab.ca/2020/04/08/preprint-recommendations/.

42. Superchi C, González JA, Solà I, Cobo E, Hren D, Boutron I. Tools used to assess the quality of peer review reports: a methodological systematic review. BMC Medical Research Methodology. 2019;19(1):48.

