## Supplementary File for "From amazing work to I beg to differ - analysis of bioRxiv preprints that received one public comment till September 2019"

### Supplementary Table 1. Description and examples of comment types left for bioRxiv preprints.

Despite public availability of comments, the wording has been modified to remove identifying information and to correct spelling errors.

| Category (major, minor) | Description | Example |
|---|---|---|
| **Author’s comment** | | |
| Publication status | Notification of submission, acceptance, update or publication of a preprint. Can also contain descriptions of changes between versions. | *“The paper has been published in the Journal of …..”* |
| Additional study information | Additional information about the study not found in the preprint, including additional analyses or links to blogs or presentations. | *“Assemblies and supplementary materials available at…”* |
| Soliciting feedback | Authors asking the community for feedback on the preprint. Can include links to other platforms where comments can be deposited. | *“We would also appreciate any comments/criticisms on the conclusions of the work.”* |
| Study promotion | Summary or a title of the preprint used as a comment. Can resemble messages, tweets or other social media status-like updates. | *“This manuscript reports our finding of a new mechanism of …”* |
| Reply or thanks for received comments | Replies or thank you message for comments received through other media (e.g. email, twitter or peer review in a journal). | *“Thanks everyone for the comments.”* |
| Other | Comments not covered by other categories (e.g. thanks to co-authors, asking where to publish the preprint, reporting misconduct…). | *“This manuscript was not approved by me or most of the other authors and may be premature.”* |
| **Non-author’s comment** | | |
| Praise | Praise or endorsement of the preprint study (or its aspects). | *“Wow, love this paper!”* |
| Suggestion | Suggesting additional methods, interpretation or results, or literature to consider. | *“I really don't want be the guy that always refer to his own research but I did publish a paper that showed…”* |
| Criticism | Criticism of the preprint content. | *“I know this is a preprint, and not the finished article - however, the reporting of … leaves a lot to be desired.”* |
| Asking for clarification | Asking for additional information or clarification on the study. | *“Do these differences correlate with sampling times of patients' cohorts?”* |
| Full peer review | Comments that stated they were peer reviews or included structure often found in journal peer review. | *“This preprint was discussed in a lab meeting and we would like to offer the following for review.”* |
| Issue detected | Issues detected with the preprint content or format (e.g. typos, missing figures or supplementary data, errors in numbers). | *“A couple of citations are incorrectly dated…”* |
| Asking for raw data or code | Asking for raw data or code that the study is based upon. | *“What about reagent availability: upon request or will you send files to…”* |
| Publication status | Asking if the version of record exists, informing if it does, or suggesting venues for publication. | *“Please, has this article been published?”* |
| Other | Comments not covered by other categories (e.g. planning to read the preprint, reporting misconduct, looking forward to other studies). | *“The downloads numbers for this preprint have been artificially inflated by a bot…”* |

Supplementary Table 2. BioRxiv subject field classification for preprints which received a single public comment between 21 May 2016 and 9 September 2019 (n=1,983).

| **Subject Area** | **Number (%)* of preprints with a single public comment from** | |
|---|---|---|
| | **non-authors** | **authors** |
| Animal Behavior and Cognition | 12 (1) | 12 (1) |
| Biochemistry | 34 (2) | 12 (1) |
| Bioengineering | 16 (1) | 10 (1) |
| Bioinformatics | 177 (9) | 76 (4) |
| Biophysics | 37 (2) | 26 (1) |
| Cancer Biology | 45 (2) | 27 (1) |
| Cell Biology | 56 (3) | 26 (1) |
| Clinical Trials | 3 (0) | 0 (0) |
| Developmental Biology | 33 (2) | 24 (1) |
| Ecology | 42 (2) | 25 (1) |
| Epidemiology | 23 (1) | 8 (0) |
| Evolutionary Biology | 108 (5) | 48 (2) |
| Genetics | 91 (5) | 30 (2) |
| Genomics | 155 (8) | 60 (3) |
| Immunology | 26 (1) | 6 (0) |
| Microbiology | 97 (5) | 45 (2) |
| Molecular Biology | 36 (2) | 26 (1) |
| Neuroscience | 230 (12) | 90 (5) |
| Paleontology | 8 (0) | 2 (0) |
| Pathology | 5 (0) | 3 (0) |
| Pharmacology and Toxicology | 9 (0) | 1 (0) |
| Physiology | 10 (1) | 6 (0) |
| Plant Biology | 40 (2) | 17 (1) |
| Scientific Communication and Education | 21 (1) | 4 (0) |
| Synthetic Biology | 12 (1) | 7 (0) |
| Systems Biology | 35 (2) | 20 (1) |
| Zoology | 5 (0) | 6 (0) |

*Percentages are calculated based on the total number of comments (n=1,983). A single field classification is chosen by the authors during preprint submission.

Supplementary Table 3. Types of single comments left by non-authors for bioRxiv preprints between 21 May 2016 and 9 September 2019 (n=1,366).

| **Comment type*** | **n (%)†** |
|---|---|
| **Praise** | **577 (42)** |
| praise and any other comment type(s) | 424 (31) |
| praise only | 86 (6) |
| praise and title or short summary of the (main) findings | 67 (5) |
| **Suggestion** | **399 (29)** |
| suggestion of literature the commenter (co-)authored | 143 (10) |
| suggesting of other literature | 118 (9) |
| **Criticism** | **226 (17)** |
| **Asking for clarification** | **213 (16)** |
| **Full peer review** | **168 (12)** |
| single person review | 87 (6) |
| group review | 81 (6) |
| **Issue detected** | **132 (10)** |
| supplementary data missing | 38 (3) |
| typo(s) | 37 (3) |
| **Asking for raw data or code** | **41 (3)** |
| **Publication status** | **34 (2)** |
| **Other** | **90 (7)** |
| research integrity concerns | 3 (0) |

*Parentages do not add up to a 100, as comment's content could contain more than one comment type. Additionally, for some categories we also present the most common subcategories. Full category coverage is defined in Supplementary Table 1.

Supplementary Table 4. Frequency of praise alongside other type of comments left by non-authors for bioRxiv preprints between 21 May 2016 and 9 September 2019 (n=1366).

| **Type of Comment*** | **Praise (n, %)** | **No praise (n, %)** |
|---|---|---|
| Suggestion | 201 (50) | 198 (50) |
| Criticism | 70 (31) | 156 (69) |
| Asking for clarification | 101 (47) | 112 (53) |
| Full peer review report | NA* | NA* |
| Issue detected | 38 (29) | 94 (71) |
| Asking for raw data or code | 18 (44) | 23 (56) |
| Publication status update | 9 (26) | 25 (74) |
| Other | 36 (40) | 54 (60) |

*We did not mark instances when praise was present in full peer review reports, as more than half were either authored by multiple authors or contained several review reports by different individuals. Of interest may be that one such review report said: "*Praises are omitted. Only concerns that may potentially improve the article are presented*".

Supplementary Table 5. Type of single comments left by authors for bioRxiv preprints between 21 May 2016 and 9 September 2019 (n=617).

| **Comment Type** | **n (%)*** |
|---|---|
| **Update on publication status** | **354 (57)** |
| link to or notification of a published version | 265 (43) |
| notification of submission to or acceptance by a journal | 62 (10) |
| description of changes between preprint and published paper | 32 (5) |
| description of changes between two preprint versions | 28 (5) |
| **Additional study information** | **158 (26)** |
| **Soliciting feedback** | **65 (11)** |
| **Self-promotion** (summary or title of the preprint) | **44 (7)** |
| **Reply or thanks for received comments** | **29 (5)** |
| reply to a comment received elsewhere | 26 (4) |
| thanks for a comment received elsewhere | 22 (4) |
| reply and sharing of received peer review comments | 5 (1) |
| **Other** | **41 (7)** |
| misconduct alert | 1 (0) |
